# Montreal Urban Observatory: research platform to monitor urban forest ecosystems for global change adaptation and health

**DOI:** 10.64898/2026.02.07.704556

**Authors:** Alain Paquette, Rita Sousa-Silva, Marine Fernandez, Maria Faticov, Laura Schillé, Emma Bacon, Elyssa Cameron, Jérémy Fraysse, Essivi Gagnon Koudji, Sarah Poirier, Jonathan Rondeau-Leclaire, Sarah Tardif, I. Tanya Handa, Isabelle Laforest-Lapointe, Danijela Puric-Mladenovic, Carly D Ziter

## Abstract

As urban populations grow, cities are increasingly seen not only as drivers of climate change but also as critical arenas for implementing mitigation and adaptation strategies. Urban forests and green infrastructure play a vital role in shaping environmental quality and human health, yet access to these benefits remains inequitable. The Montreal Urban Observatory was designed to investigate the complex relationships among biodiversity, ecosystem functioning, and human health. Twenty-five permanent plots were established across the Island of Montreal, strategically located along gradients in vegetation cover, population density, and mean household income. Interdisciplinary research within the Observatory focuses on themes including urban forest structure and function, abiotic conditions, biodiversity, ecological interactions, nature’s contributions to people, related human health outcomes, and developing novel methodological approaches. The Observatory supports long-term, cross-scale monitoring and fosters collaborative research on urban social-ecological systems, thus contributing to global initiatives to enhance urban sustainability and equity through nature-based solutions. Early results confirm the relevance of the main gradients for several organisms and response variables, from microbes to trees and abiotic factors. For example, sampling all trees, public and private, within a radius of 200m centered on each plot revealed significant differences in the diversity and structure of private and public trees, including an overwhelming dominance of *Thuya occidentalis* not captured in commonly used public tree inventories.

## Introduction

In an era of rapid urbanization, cities have become focal points for responding to environmental and societal challenges. Over 70% of the world’s population is projected to live in cities in the next 30 years (Lwasa et al. 2022). This growing urbanization has a significant impact on biodiversity (Barnosky et al. 2011, Isbell et al. 2015), with substantial consequences for human health and well-being (Pascual et al. 2017, Bratman et al. 2019). As extreme heat, unpredictable weather events, and unexpected challenges such as the recent COVID-19 pandemic become more common (Mahadevia et al. 2025), researchers and policymakers are increasingly turning to nature-based solutions within a “One Health” approach (Romaneli and Taberner 2024, WHO 2025). One key strategy is the expansion of urban green infrastructure, aimed at creating resilient and sustainable cities (Lafortezza and Sanesi 2019).

Around the world, urban green infrastructure, including parks, gardens, street trees and green roofs, plays a critical role in determining nature’s contributions to people. These contributions span a range of ecosystem services, such as climate regulation (Emmanuel and Loconsole 2015, Marando et al. 2022), air purification (Vieira et al. 2018), managing stormwater (Livesley et al. 2016) and physical and mental health benefits (Donovan et al. 2018, Turner□Skoff and Cavender 2019). For example, proximity to green spaces is associated with stress reduction, improved mental health, and opportunities for recreation (Sandifer et al. 2015, Wood et al. 2017). Among green infrastructure components, urban forests, defined as all trees across public and private urban spaces, are particularly important. They not only help regulate urban microclimate, sequester carbon and provide habitat for city wildlife, but also foster everyday human–nature interactions (Soga and Gaston 2020, Moreira et al. 2025). While benefits of urban forests are substantial, efforts to expand them should also consider ecosystem disservices, such as allergies of urban residents to tree pollen whose seasonality and drivers are poorly understood (Sousa-Silva et al. 2020, Sousa-Silva et al. 2021). Another concern is the spread of zoonoses such as Lyme disease, West Nile virus, and other tick- or mosquito-borne diseases, which are increasingly linked to urbanization and climate change. For instance, poorly maintained or ill-designed urban forests can create breeding habitats for mosquito vectors, with implications for disease transmission in both temperate and tropical urban settings (Leighton et al. 2012, Mathieu and Karmali 2016, Guo et al. 2022, Fournet et al. 2024). Overall, a deeper understanding of how urban forests contribute to both ecosystem services and disservices is essential for evidence-based planning and equitable implementation of nature-based solutions in cities (Cook et al. 2025).

While the ecological and social value of urban forests is well recognized, cities continue to face significant challenges in implementing inclusive and evidence-based urban forest planning, management and stewardship (Pataki et al. 2011, Sousa-Silva et al. 2023a). These challenges stem in part from a limited understanding of the relationships among urban form, urban forests, biodiversity, and human well-being at the appropriate spatial and temporal scales (Sandifer et al. 2015). For example, where, and how many trees of what types are needed to fulfill conservation, recreational and well-being purposes? What are the strengths and shapes of these relationships? Are they consistent across urban areas that differ in their built density and design? It is crucial to address these questions in light of ongoing climate change, which is already causing significant adverse effects on urban communities facing systemic environmental inequities. Unfortunately, equity-deserving communities are also the least likely to have access to high tree cover and biodiversity (Schell et al. 2020) and are thus more likely to develop health problems following environmental exposure (e.g., to noise, pollution, heat). This calls for urgent mitigation and adaptation efforts. Interdisciplinary research is therefore needed, not only to monitor and maintain or increase biodiversity in urban areas, but also to inform planning and management strategies that ensure equal access to nature’s benefits (Frantzeskaki et al. 2025, Ziter and Buxton 2025).

To better anticipate and manage both the benefits and disservices of urban forests and associated green infrastructure, particularly in relation to human health and well-being, research must be conducted at the appropriate spatial scales (Uchida et al. 2021). Interventions that work in one neighbourhood will not necessarily hold in another, particularly given differences in urban form and socio-demographics. Understanding the mechanisms underlying nature’s contributions to people is an important frontier in ecological and public health research (Sandifer et al. 2015). This requires characterizing the strength and direction of associations among multiple indicators of biodiversity and multiple (dis)services across the urban landscape and tracking these changes over time. Monitoring and establishment of permanent sampling plots have provided much needed insight into similar questions across other systems. For example, globally distributed experimental networks, where standard data are collected across gradients of environmental conditions and over time, have provided substantial insight into the generality and site-specificity of factors influencing biodiversity and ecosystem functioning of grassland (Borer et al. 2014, Lind 2016) and forest systems (Paquette et al. 2018). However, such monitoring have been slower to develop in cities, which are challenging due to the high spatial heterogeneity, temporal dynamism, complex governance and land management structures (Aronson et al. 2017, Dyson et al. 2019). While local urban observatory networks have been established, such as the CityScapeLab Berlin (von der Lippe et al. 2020), such efforts still remain limited compared to non-urban systems like the National Ecological Observatory Network (NEON) (Schimel et al. 2007).

USA’s Long-Term Ecological Research (LTER) network launched in 1997 its first urban sites: Baltimore Ecosystem Study and Central Arizona – Phoenix. They pioneered urban system science and produced some of the most influential research on urban watersheds, biodiversity, and people’s relationships to nature (Bang et al. 2012, Grove et al. 2015, Groffman et al. 2017, Pickett et al. 2017). In Ontario, Canada, in-depth monitoring and observation of urban forests have been conducted, for over three decades, through two initiatives. The Neighbourwoods© (Kenney and Puric-Mladenovic 1995) and Vegetation Sampling Protocol programs (Puric-Mladenovic 2015) have emphasized systematic monitoring and the evaluation of the structure and functionality of urban trees across a variety of climatic conditions, land uses, and timeframes in Southern Ontario, with many thousand plots. Data collected about trees, ranging from a single land parcel to an entire town, have provided essential insights into the structure, composition, and function of urban forests, and supported studies on carbon sequestration and others (Drever et al. 2021). Much of the existing sampling thus covers urban woodlots and watersheds, but does not focus on trees in streets, parks, and especially on private land. Building on these ideas, needs, and prior initiatives, two large plots were installed in Montreal, which documented strong differences in tree diversity and abundances between public and privately owned lands, as well as among the different private land uses (Hutt-Taylor and Ziter 2022, Sousa-Silva et al. 2023b). Broadening this approach to create a network of plots that captures the diversity of the urban environment is essential for understanding the generality of these findings and testing the underlying mechanisms.

Here we introduce the Montreal Urban Observatory, a network of 25 permanent urban plots that has the potential to answer not only specific research questions but also larger, interconnected ones, with an interdisciplinary One Health approach. By analyzing data from these plots, we aim to enhance our understanding of the mechanisms that connect ecosystem functioning, benefits, and public health. Additionally, we seek to provide a model for studying these processes in cities around the world, which will help clarify which relationships are generalizable and which are context specific.

### Network implementation

The Montreal Urban Observatory comprises 25 plots located across the Island of Montreal (Fig. 1; Table S1). The island, which includes the City of Montreal and 15 other municipalities, covers an area of 500 km^2^ and has a population of 2.2 million inhabitants (Institut de la statistique du Québec 2024). Plots were established in 2021 across the Island to represent a diversity of contexts based on three gradients: vegetation density, population density and mean household income. Sites were not chosen to be proportionally representative of land cover or socio-demographics, but rather to capture variation of multiple variables hypothesized to drive biodiversity and ecosystem function in urban systems. This variability improves the capacity for landscape-scale inference, including the detection of non-linear relationships (Eigenbrod et al. 2011, Miller et al. 2021), and provides a template for future cross-city comparisons.

**Figure 1.**
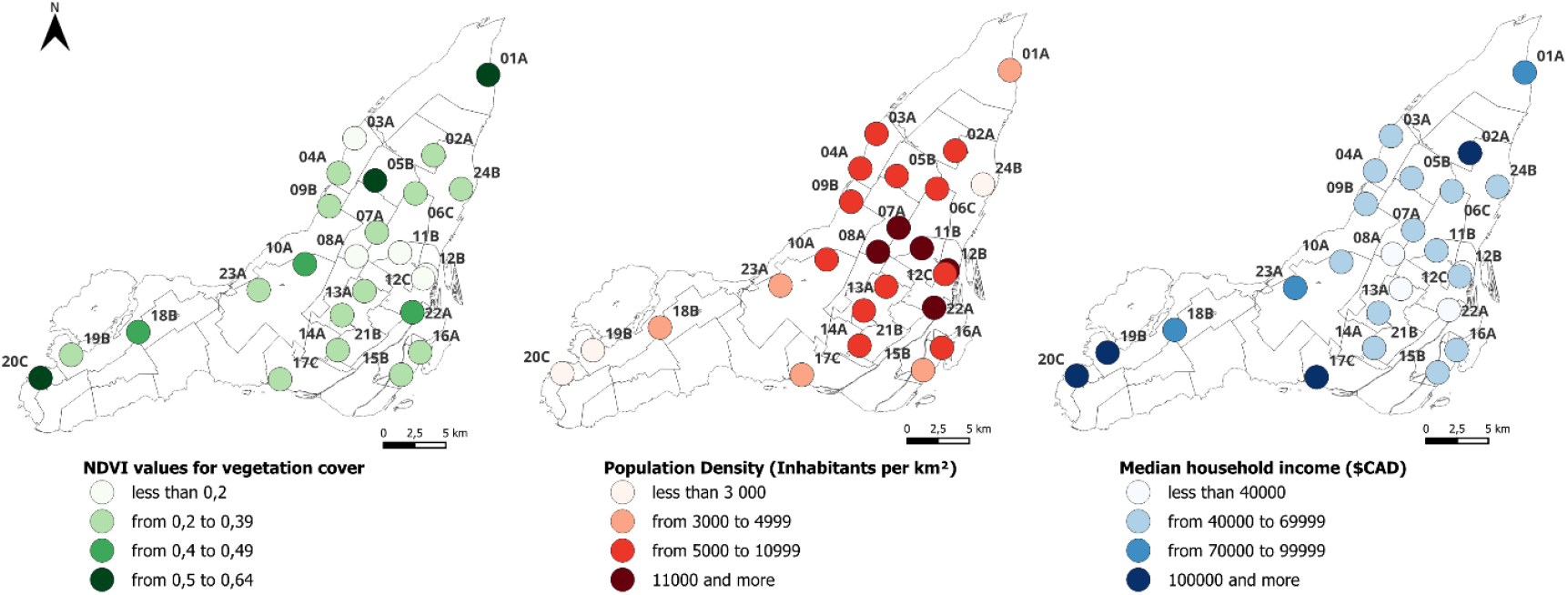
Plot locations on the Island of Montreal along the three distinct gradients.

We began with vegetation cover as the primary gradient, as it directly reflects the structure of the urban forest and vegetation, which is hypothesized to underpin both ecological and social benefits. Vegetation data were obtained from the Canadian Urban Environmental Health Research Consortium (Gorelick et al. 2017). NDVI values (2019) were assigned to six-character postal codes and interpolated to a 30 m grid cell using inverse distance weighting. The resulting vegetation index was scaled from 0 to 100%. The second gradient, population density, was used to represent variations in the distribution and concentration of residential populations across the study area. When combined with vegetation data, it helps identify areas where residents are more likely to be exposed to urban greenery. Population data were obtained from Statistics Canada’s DMTI 2019 dataset (DMTI Spatial Inc), aggregated at the forward sortation area level (i.e., the first three digits of a postal code), and interpolated to a 30 m grid. Each grid cell was assigned a relative density index based on the ratio of local to maximum population density on the Island of Montreal.

The vegetation and population density indices were combined by addition into a composite index ranging from 0 to 200%. The resulting map was smoothed using focal mean statistics with a 1000 m radius, and values were grouped into ten percentile-based categories. To prioritize areas with both vegetation and human presence, we retained only the five highest categories of the composite index. We then selected candidate locations within these areas, applying a minimum spacing of 500 m between points and an exclusion buffer around each point to avoid overrepresentation. The exclusion radius ranged from 1 km to 2.5 km, with larger buffers in lower-scoring areas to increase the likelihood of placing plots in areas with more vegetation and higher population density. We removed overlapping candidate plots and prioritized more central locations when multiple candidate plots were available, particularly in the east and west of the Island of Montreal. This process resulted in a selection of 21 plots distributed across the combined vegetation-population gradient.

To ensure the network also captured socio-economic variation, we supplemented the initial network of 21 plots with four additional plots selected based on household income. We used median household income data from the 2016 Canadian Census at the dissemination area level. These additional plots were placed in areas underrepresented in the initial selection but important for capturing income-related disparities, bringing the total number of plots to 25. Note that the three gradients were constructed only for the purpose of establishing the network in 2021, using the most accurate and recent data available at the time. Updated values should be used for the purpose of analyzing their relationships with field observations.

To understand the composition and structure of the urban forest across the network, we conducted a complete tree inventory in the summers of 2023 and 2024. We identified and measured diameter at breast height (DBH) for all accessible trees within a 200 m radius of the plot center. All trees were geolocated and identified to the species level when possible (97%). For multi-stemmed trees, all stem measurements were consolidated into one individual DBH as per Magarik et al. (2020). We used Montreal’s public tree database (donnees.montreal.ca) to locate all publicly managed trees. In some boroughs and municipalities, tree inventory data were not available. In these cases, we considered all trees on land owned or managed by the city or municipality to be publicly owned trees (e.g., trees found on the public right-of-way and in parks) and those on land not managed by the city or municipality to be privately owned (e.g., residential yards, commercial land, educational or faith-based institutions). To access trees on private land, we obtained permission from landowners or estimated species and DBH from the street when visible. We inventoried a total of 34,370 trees spanning 101 genera and 320 species, from seedlings with a measurable DBH to large trees up to 175 cm in diameter (Table 1). The three most common species were Eastern white cedar (*Thuja occidentalis*), found almost exclusively as residential hedges, Norway maple (*Acer platanoides*), and common lilac (*Syringa vulgaris*), representing 42%, 6%, and 4% of all trees across the observatory, respectively. Whereas Norway maple is a very common urban tree in northeastern North-America (Nock et al. 2013), the other two species, common lilac and particularly cedar, were unexpected as they are never reported as important in public inventories. This difference in tree species composition has implications for ecosystem function and services (Hutt-Taylor and Ziter 2022, Sousa-Silva et al. 2023b), and is a strong demonstration of the importance of more extensive sampling. Indeed, most of the trees inventoried were on private land (71%), showing that in the areas representative of where most people live, the proportion of trees on private land can be significantly higher than the often cited 50%.

**Table 1.**
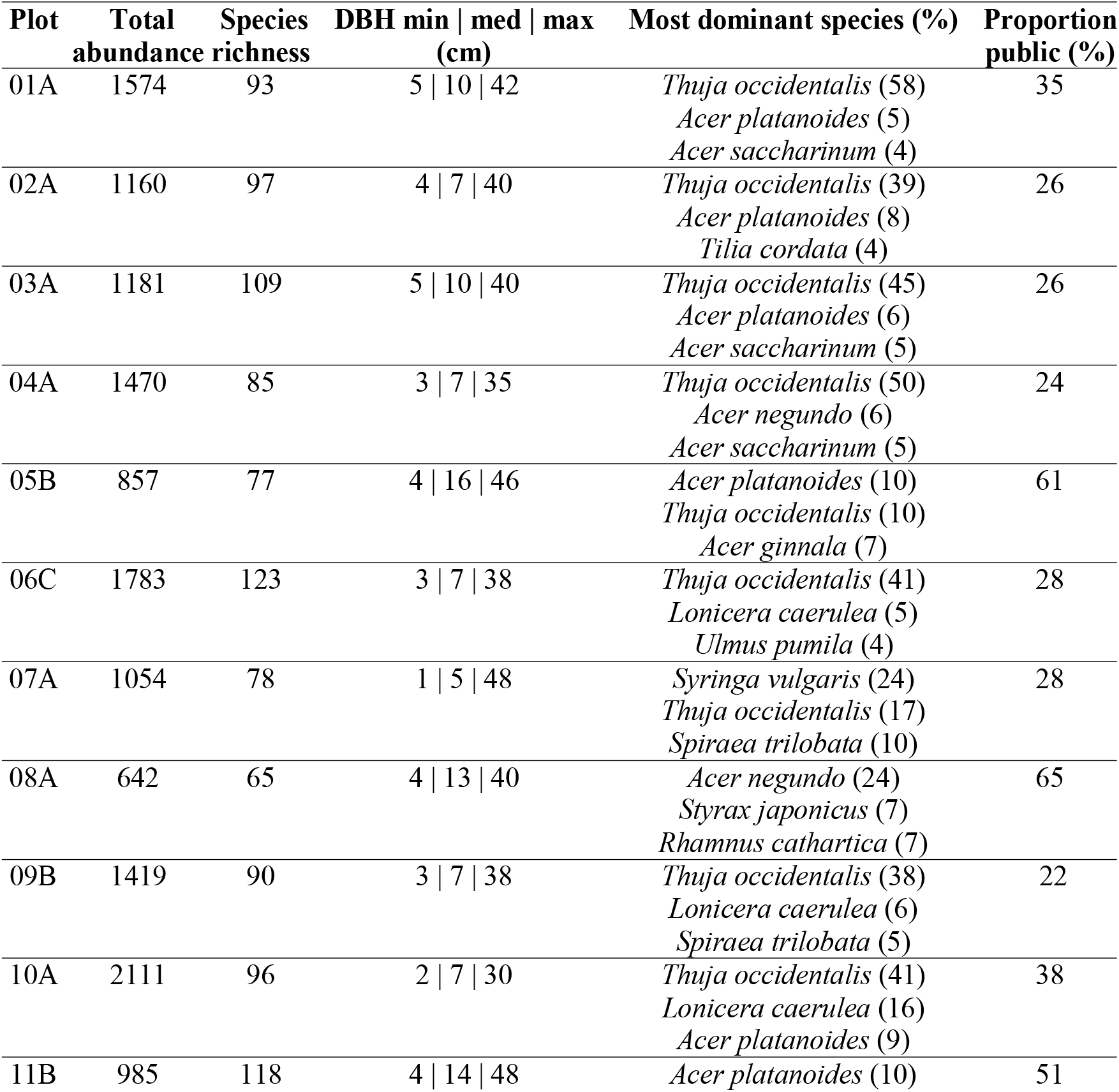
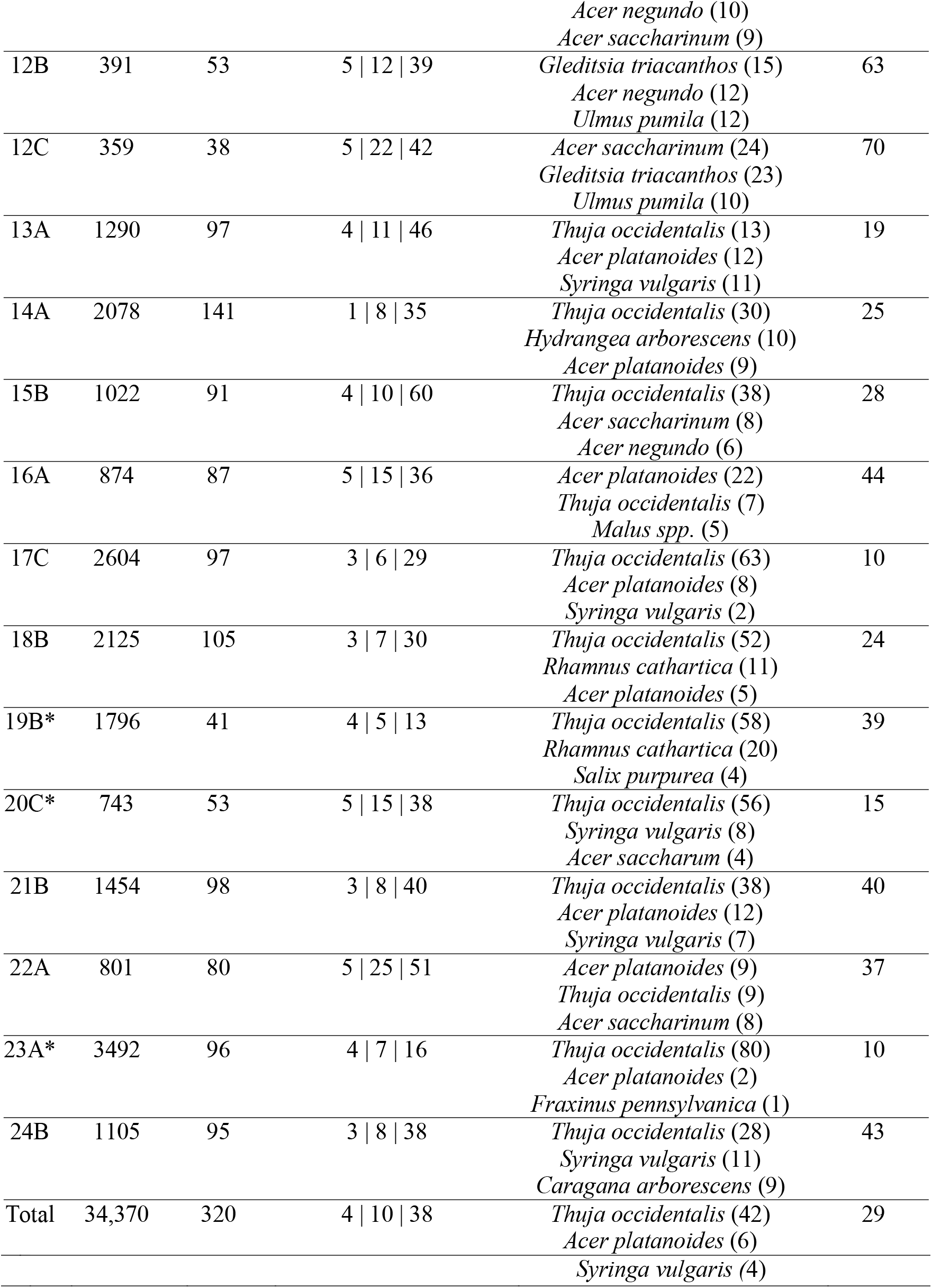
Summary statistics for the tree inventories. Values represent tree abundance, species richness, the 10^th^ and 90^th^ percentiles minimum, median, and maximum diameter at breast height (DBH; cm), for each plot. Most dominant species are also provided with their relative abundance in parentheses, as well as the proportion of all trees that is on public land. Public land included publicly managed green spaces such as public right-of-way and parks. Private land is any land use type that is not publicly managed, including residential, commercial, and institutional land. *Plots that included large, densely forested areas that were sampled using classic forestry techniques but are not included here.

Since the network was not designed to be representative of the entire city but rather to maximize coverage over gradients of interest, plot-level data cannot be directly extrapolated without adjustment. However, if city-wide inferences are preferable, this can be achieved by applying correction factors (or multipliers), based on proportion representation for each gradient of interest (see Table S2 for an example).

### Overview of projects and objectives, and early results

The Montreal Urban Observatory allows for the monitoring of biodiversity and its interactions with the urban environment. To this end, several projects are currently being carried out, which are organized into seven interconnected themes, reflecting the diverse and multifaceted nature of urban research (Fig. 2).

1. **Structure and function of the urban forest:** This theme encompasses studies on the composition and dynamics of urban trees, including inventories of public and private trees, stakeholder-driven reflections on species selection and planting locations, and research on tree ecophysiology to understand how urban conditions influence tree health and performance.
2. **Abiotic environment:** Projects within this theme focus on the physical and chemical properties of the urban environment, such as soil characteristics (temperature, humidity, compaction, and organic matter, carbon, nitrogen, phosphorus and calcium content, among others), ground vegetation cover, and air quality.
3. **Biodiversity:** This theme covers multi-taxa biodiversity assessments, including inventories of trees, birds, tree-associated arthropods, flying insects, and soil fauna, contributing to a clearer picture of urban biodiversity across multiple taxonomic groups. For example, ongoing work is examining the impacts of urbanization and tree diversity on avian diversity and insectivory (Schillé et al. 2025).
4. **Ecological and trophic interactions**: Research in this theme explores the intricate web of interactions within urban ecosystems. Examples include studies on root and leaf microbiomes, herbivory patterns, and trophic interactions between insectivorous birds and herbivorous insects.
5. **Nature’s contributions to people:** Projects here evaluate the services and disservices provided by urban forests. Examples include the role of urban trees in mitigating urban heat islands, providing microhabitats, and regulating pest insect populations, alongside potential disservices such as allergenic pollen production.
6. **Human health and well-being**: This theme addresses the connections between urban forests and human health, including for example studies on pollen distribution patterns and emergency admission for asthma to better understand how urban vegetation influences respiratory health, among other outcomes related to physical and mental well-being. Under a One Health approach, this theme also incorporates equity dimension by examining how exposure to urban forests and environmental stressors varies across neighborhoods with different socio-economic profiles.
7. **Innovative methods and technologies**: Our observatory embraces cutting-edge methodologies to advance urban ecology research. Examples include acoustic ecology for bird community monitoring using autonomous recording units, custom-built high-speed cameras for capturing trophic interactions, DNA metabarcoding for insect sampling, and flow cytometry for detailed pollen analysis.

**Figure 2.**
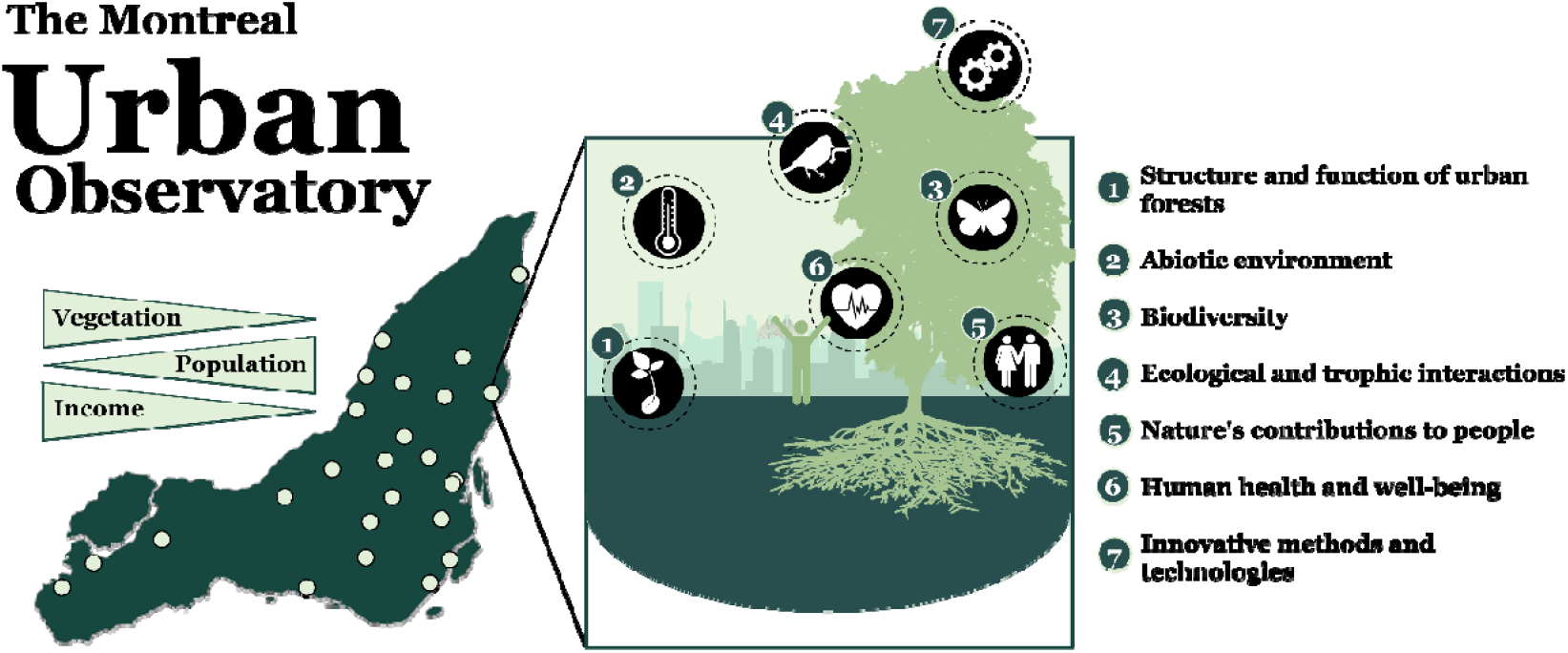
The seven interconnected research themes of the Observatory, reflecting the diverse and multifaceted nature of urban research.

**Figure 3.**
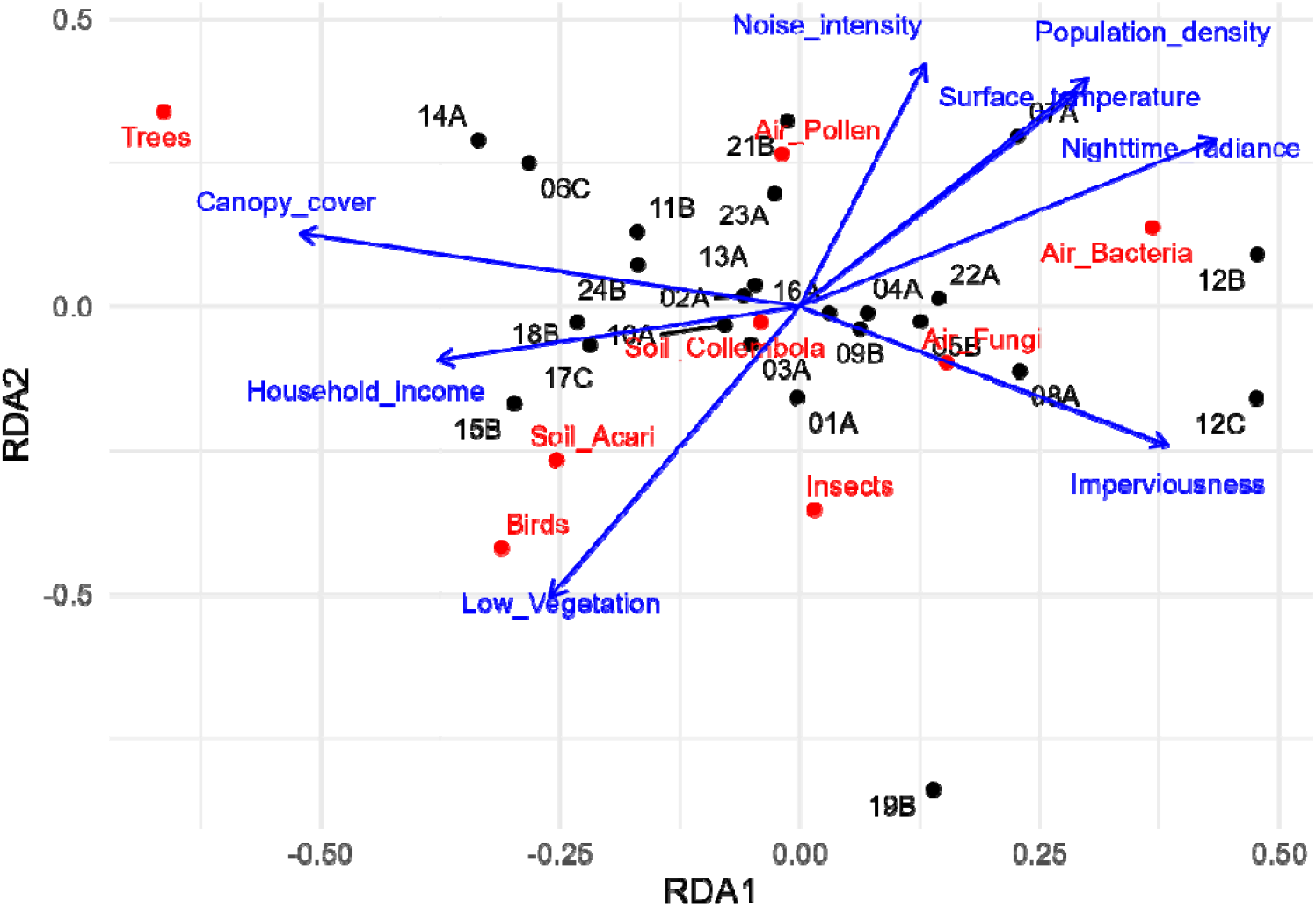
Redundancy analysis (RDA) showing the relationships among study plots (black labels), explanatory environmental and socio-economic variables (blue arrows; Table S4), and the richness of different groups of organisms sampled (red labels; Table S3). Axis scaling is based on the RDA ordination; arrow directions indicate the correlation of variables with the ordination axes.

To illustrate the structuring of the Urban Observatory of Montreal along the urbanization gradient, we performed a redundancy analysis (RDA; vegan package, R version 4.5.0) of the position of study plots along the urban gradient based on the specific richness of several taxonomic groups (Table S3). The response matrix was not composed, as usual, of the abundances of a given taxon, but rather of the specific richness of eight groups of organisms already sampled in the Urban Observatory: birds, insects, trees, air bacteria, air fungi, air pollen, soil acari, and soil collembola. We transformed these richness data using the Hellinger method to reduce the effect of scale differences. The explanatory matrix consisted of eight predictors reflecting the multidimensional nature of urbanization (Table S4): canopy (> 3m) and low (< 3 m) vegetation cover, population density, average household income, impervious surface cover, nighttime radiance as a proxy for light pollution, sound intensity as a proxy for noise pollution, and surface temperature as a proxy for the urban heat island effect (Table S5 for details). All environmental variables were standardized (centered and scaled) to account for differences in units and magnitude. A spatial aggregation level of 200 m around the center of each plot was used for all predictors.

The RDA thus provided a simultaneous representation of the relationships among sites, taxonomic groups, and environmental variables. Our plots do spread nicely along the main gradients, as expected. We find that low-income neighborhoods have greater population density, higher surface temperatures, more noise and light pollution. Plots with greater canopy cover and lesser imperviousness have the most tree species richness. Insects, birds and Acari, for their part, seem to respond better to low vegetation in high income neighbourhoods, while other groups of organisms such as Fungi and Collembola do not seem to vary much within our network. This could be explained by the scale at which they were sampled, and finer-scale analyses are of course possible because all samples such as trees are geo-localized.

We are confident the Observatory will continue to support several projects in the future, with the aim of responding to ecological issues and highlighting innovative solutions linked to the understudied urban environment. Accordingly, we extend an open invitation to researchers, practitioners, policymakers, and community stakeholders to collaborate with us in using the Montreal Urban Observatory towards advancing our understanding of urban forests and their critical role in our shared environment and help to establish parallel urban networks in cities across the globe. By combining and expanding efforts and knowledge, we can work towards ensuring the long-term health, resilience, and sustainability of these essential ecosystems. Together, we can build a future where urban forests are more equitably distributed, thriving and supporting both ecological and societal well-being for generations to come.

## Acknowledgements

We thank Daniel Lesieur and Mélanie Desrochers (CEF professionals), as well as Hendrick Paquette Ambroise (UQAM intern) for their important contribution to the establishment of the Montreal Observatory. NDVI metrics, indexed to DMTI Spatial Inc. postal codes, were provided by CANUE (Canadian Urban Environmental Health Research Consortium) and the Google Earth Engine Team, using Landsat 5 and 8 TM Annual Greenest-Pixel TOA Reflectance Composite. We also recognize the financial support of two Natural Sciences and Engineering Research Council of Canada (NSERC) programs: Alliance (AP) and New Frontiers in Research (ILL and AP) and that of the Fonds vert dans le cadre du Plan d’action 2013-2020 sur les changements climatiques du gouvernement québécois.

