## Supplementary Tables S1 to S5 for "Montreal Urban Observatory: research platform to monitor urban forest ecosystems for global change adaptation and health"

Table S1. Location of the 25 urban permanent plots in the Montreal Urban Observatory Network.

| Plot ID | Longitude | Latitude |
|---|---|---|
| 01A | -73.4934 | 45.6621 |
| 02A | -73.5502 | 45.6039 |
| 03A | -73.6309 | 45.6161 |
| 04A | -73.6469 | 45.5909 |
| 05B | -73.6099 | 45.5856 |
| 06C | -73.5684 | 45.5764 |
| 07A | -73.6083 | 45.5484 |
| 08A | -73.6283 | 45.5315 |
| 09B | -73.6571 | 45.5671 |
| 10A | -73.6817 | 45.5254 |
| 11B | -73.584 | 45.5333 |
| 12B | -73.5572 | 45.5174 |
| 12C | -73.5603 | 45.5148 |
| 13A | -73.6209 | 45.5058 |
| 14A | -73.6473 | 45.4628 |
| 15B | -73.5833 | 45.4453 |
| 16A | -73.5629 | 45.4614 |
| 17C | -73.7071 | 45.4418 |
| 18B | -73.853 | 45.4755 |
| 19B | -73.9217 | 45.4589 |
| 20C | -73.9537 | 45.4419 |

| Plot ID | Longitude | Latitude |
| --- | --- | --- |
| 21B | -73.6434 | 45.4882 |
| 22A | -73.5713 | 45.4904 |
| 23A | -73.7294 | 45.5064 |
| 24B | -73.5212 | 45.5795 |

Table S2. Example of correction factors (or multipliers) that could be used to make city-wide inference based on the true proportional representation of our plots, here for population density. The study area is first divided into ten percentile-based classes (for a given gradient) and the proportion of the city represented by each class is then calculated, which can be used as correction factor. Population density data for this table were obtained from Statistics Canada's DMTI 2019 dataset, aggregated at the forward sortation area (FSA) level.

| **Plot ID** | **Class** | **Area (km$^{2)}$** | **Proportion** |
|---|---|---|---|
| 01A | 2 | 107.43 | 0.216 |
| 02A | 4 | 72.29 | 0.145 |
| 03A | 4 | 72.29 | 0.145 |
| 04A | 5 | 29.52 | 0.059 |
| 05B | 4 | 72.29 | 0.145 |
| 06C | 5 | 29.52 | 0.059 |
| 07A | 8 | 5.98 | 0.012 |
| 08A | 10 | 1.66 | 0.003 |
| 09B | 5 | 29.52 | 0.059 |
| 10A | 4 | 72.29 | 0.145 |
| 11B | 9 | 1.94 | 0.004 |
| 12B | 6 | 18.79 | 0.038 |
| 12C | 6 | 18.79 | 0.038 |
| 13A | 7 | 16.62 | 0.033 |
| 14A | 5 | 29.52 | 0.059 |
| 15B | 4 | 72.29 | 0.145 |
| 16A | 7 | 16.62 | 0.033 |
| 17C | 3 | 81.11 | 0.163 |
| 18B | 3 | 81.11 | 0.163 |
| 19B | 1 | 162.92 | 0.327 |
| 20C | 1 | 162.92 | 0.327 |
| 21B | 4 | 72.29 | 0.145 |
| 22A | 6 | 18.79 | 0.038 |
| 23A | 1 | 162.92 | 0.327 |
| 24B | 2 | 107.43 | 0.216 |

Table S3. Per plot richness of several groups of organisms (at the species, genus, or molecular sequence level depending on group; see note).

| Plot ID | Birds | Pollen | Bacteria | Fungi | Acari | Collembola | Trees | Insects |
|---|---|---|---|---|---|---|---|---|
| 01A | 14 | 18 | 344 | 183 | 52 | 18 | 93 | 16 |
| 02A | 14 | 28 | 373 | 166 | 70 | 23 | 97 | 6 |
| 03A | 19 | 32 | 408 | 201 | 66 | 32 | 109 | 12 |
| 04A | 10 | 26 | 374 | 161 | 63 | 26 | 85 | 10 |
| 05B | 12 | 30 | 359 | 171 | 51 | 31 | 77 | 10 |
| 06C | 8 | 34 | 317 | 134 | 57 | 23 | 123 | 10 |
| 07A | 3 | 41 | 349 | 167 | 45 | 20 | 78 | 8 |
| 08A | 9 | 27 | 383 | 152 | 64 | 29 | 65 | 13 |
| 09B | 11 | 32 | 365 | 208 | 61 | 34 | 90 | 11 |
| 10A | 12 | 21 | 338 | 159 | 57 | 33 | 96 | 10 |
| 11B | 7 | 33 | 350 | 160 | 70 | 25 | 118 | 13 |
| 12B | 4 | 27 | 367 | 157 | 32 | 48 | 53 | 11 |
| 12C | 4 | 37 | 355 | 162 | 73 | 22 | 38 | 12 |
| 13A | 9 | 26 | 340 | 169 | 58 | 36 | 97 | 10 |
| 14A | 8 | 38 | 320 | 151 | 53 | 33 | 141 | 12 |
| 15B | 12 | 36 | 318 | 135 | 126 | 47 | 91 | 7 |
| 16A | 15 | 33 | 352 | 183 | 50 | 37 | 87 | 9 |
| 17C | 15 | 41 | 318 | 159 | 79 | 39 | 97 | 10 |
| 18B | 15 | 39 | 350 | 145 | 85 | 38 | 105 | 10 |
| 19B | 24 | 18 | 303 | 176 | 90 | 25 | 41 | 24 |
| 20C | NA | 19 | 328 | 85 | 32 | 12 | 53 | 18 |
| 21B | 8 | 39 | 384 | 183 | 78 | 41 | 98 | 0 |
| 22A | 10 | 41 | 382 | 187 | 62 | 29 | 80 | 11 |
| 23A | 11 | 47 | 343 | 144 | 41 | 30 | 96 | 13 |
| 24B | 9 | 31 | 299 | 136 | 56 | 47 | 95 | 10 |

To sample bird diversity, an AudioMoth (Hill et al. 2018) device was placed in June 2023 in the center of each plot to passively record bird vocalizations during the morning chorus. From these recordings, four 10-minute samples per plot were selected for species identification by an expert ornithologist (Schillé et al. 2025).

Bacterial, fungal and plant taxonomic diversity of bioaerosols was assessed by amplicon sequencing of the 16S, ITS and trnL genes, respectively. Samples were collected by concentrating air using an aerosol collector (SASS®4100, Research International) over three sampling rushes in 2022 (May 3rd-7th, June 8th-13th, and August 31st-September 9th) for a total of 65 samples. Soil arthropods (Acari and Collembola) were assessed by amplicon sequencing of the 18S gene. Soil samples were collected in July 2022 under three tree individuals of each of two tree species (*Acer platanoides* and *Acer saccharinum*) at all 25 sites yielding a total of 150 samples. Soils were cored (diameter = 5.08 cm) to a depth of 10 cm within a 1 m radius of each tree and tools were sterilized between samples (10% bleach). Soils were sieved (2 mm), homogenized, transported at 4°C, and stored for DNA extraction at -80°C. Before DNA amplification, a pre-lysis step was performed by mixing soil with freshly prepared phosphate buffer (pH 8; $Na_2HPO_4$ 0.12 M) for 15–30 min, followed by centrifugation (10 000 rcf, 10 min) (Taberlet et al. 2012). The supernatant containing extracellular DNA was extracted with the PowerSoil kit (Qiagen., Germany). We used a two-primer cocktail for amplification of the 18S and COI genes, but only 18S are used here. PCR conditions followed Kirse et al. (2021) with a 55 °C annealing temperature.

DNA for all taxa was sequenced on Illumina Miseq PE300 (Illumina, San Diego, CA, USA). Reads were resolved to amplicon sequence variants (ASVs) using the DADA2 pipeline, processing each amplicon dataset separately. Sequence tables were rarefied to the smallest sample size per dataset to remove detection bias from uneven sample sequencing depth. For arthropods, taxonomic assignment to Acari and Collembola was done with the SILVA library database. Richness for all taxa is reported as the number of ASVs (i.-e., the number of genetic variants) per sample.

All trees, public and private, were identified to species and their diameter at breast height (DBH; 1,3m) measured over the whole plots (200m radius). See main text for more on trees.

Insects were collected twice during June 2023 from a series of pan traps (Moericke 1951) operated for 48 hours during each sampling at each site. Insects were stored in the freezer at -20°C in 15 mL Falcon tubes. They were later visually assessed at the family-level taxonomic richness using a Leica M165 FC stereo microscope. The data in the table represent the cumulative family richness of the two sampling periods.

Table S4. Per plot values for key environmental and socio-economic variables that are linked to urbanization gradients and hypothesized to affect urban biodiversity. See Table S5 for details.

| Plot | Canopy cover (%) | Low Vegetation (%) | Population density (people km$^{-2}$) | Household income ($ CAD) | Imperviousness (%) | Nighttime radiance (nW cm$^{-2}$ sr$^{-1}$) | Noise intensity (dB) | Surface temperature (°C) |
|---|---|---|---|---|---|---|---|---|
| 01A | 32.61 | 29.75 | 3720 | 38076 | 43.23 | 25.37 | 62.49 | 30.80 |
| 02A | 14.23 | 25.71 | 10920 | 33758 | 47.65 | 43.25 | 52.58 | 36.38 |
| 03A | 15.97 | 24.22 | 4393 | 31347 | 41.18 | 31.04 | 51.05 | 32.92 |
| 04A | 17.58 | 12.84 | 10020 | 32709 | 44.90 | 31.62 | 62.10 | 33.35 |
| 05B | 18.31 | 32.27 | 7943 | 31632 | 52.19 | 34.45 | 50.31 | 32.07 |
| 06C | 23.28 | 16.81 | 10561 | 37273 | 49.06 | 43.95 | 62.15 | 36.80 |
| 07A | 14.18 | 9.53 | 18123 | 29992 | 52.59 | 37.96 | 65.54 | 38.87 |
| 08A | 10.34 | 9.98 | 17831 | 25683 | 48.51 | 69.74 | 53.91 | 38.87 |
| 09B | 19.91 | 13.43 | 10060 | 34771 | 41.06 | 40.67 | 53.10 | 32.24 |
| 10A | 21.03 | 26.26 | 5437 | 30843 | 43.29 | 34.62 | 53.82 | 31.71 |
| 11B | 26.58 | 5.59 | 15723 | 42980 | 55.41 | 41.71 | 56.82 | 37.25 |
| 12B | 8.87 | 1.28 | 17174 | 31354 | 69.74 | 75.15 | 58.70 | 41.13 |
| 12C | 8.59 | 2.71 | 9443 | 34168 | 60.92 | 103.77 | 62.22 | 41.08 |
| 13A | 28.08 | 8.67 | 9749 | 34978 | 48.00 | 40.52 | 65.14 | 35.06 |
| 14A | 25.42 | 34.12 | 8380 | 33741 | 50.88 | 35.41 | 52.94 | 31.58 |
| 15B | 23.73 | 22.75 | 11405 | 37966 | 51.62 | 23.58 | 55.30 | 35.02 |
| 16A | 16.60 | 17.54 | 10051 | 37722 | 51.29 | 32.08 | 61.63 | 39.96 |
| 17C | 30.17 | 24.83 | 5273 | 44395 | 41.94 | 20.84 | 56.46 | 34.59 |
| 18B | 21.59 | 30.56 | 3284 | 37565 | 44.87 | 18.50 | 50.64 | 32.82 |
| 19B | 15.28 | 64.99 | 375 | 38295 | 68.71 | 5.29 | 44.12 | 27.24 |
| 20C | 73.13 | 22.00 | 74 | 45600 | 24.35 | 2.99 | NA | 22.13 |
| 21B | 23.73 | 11.67 | 13110 | 29834 | 51.65 | 68.65 | 62.02 | 36.77 |
| 22A | 28.81 | 11.36 | 15708 | 27325 | 43.30 | 73.87 | 61.16 | 35.32 |
| 23A | 17.10 | 26.08 | 4857 | 45486 | 46.96 | 54.07 | 52.02 | 33.06 |
| 24B | 18.72 | 15.94 | 4369 | 37842 | 52.78 | 70.62 | 61.32 | 39.59 |

Table S5. Methodological details for the key environmental and socio-economic variables presented in Table S4.

| Variable | Unit | Methodology | Data source | Resolution | Aggregation level |
|---|---|---|---|---|---|
| **Canopy cover** | % | Estimated by combining the 2023 Normalized Difference Vegetation Index (NDVI) with the digital elevation model (DEM) of the city. Areas with NDVI ≥ 0.3 and DEM ≥ 3 m were classified as canopy cover. | (Communauté métropolitaine de Montréal 2023) | 1m$^2$ | 200m around plot center. |
| **Low vegetation cover** | % | Areas with NDVI ≥ 0.3 and DEM< 3 m were classified as low vegetation cover. | (Communauté métropolitaine de Montréal 2023) | 1m$^2$ | 200m around plot center. |
| **Population density** | people/km$^2$ | Extracted from the 2021 Census and reported for each Dissemination Area (DA) as defined by Statistics Canada. | (Statistics Canada 2021) | Dissemination Area (DA) | Weighted mean based on the area of each DA and the corresponding population density falling within a 200-m buffer around plot center. |
| **Household income** | $ CAD | Extracted from the 2021 Census and reported for each Dissemination Area (DA) as defined by Statistics Canada. | (Statistics Canada 2021) | Dissemination Area (DA) | Weighted mean based on the area of each DA and the corresponding population density falling within a 200-m buffer around plot center. |
| **Impervious cover** | % | Areas with NDVI < 0.3 were classified as impervious cover. | (Communauté métropolitaine de Montréal 2023) | 1m$^2$ | 200m around plot center. |
| **Nighttime radiance** | nW/cm$^2$/sr | Annual cloud-free Day/Night Band composite data derived from 2023 NOAA VIIRS DNB observations, reprocessed by the Earth Observation Group (EOG) of the National Geophysical Data Center to provide annual data. | (Elvidge et al. 2013) | 450 m spatial resolution | Area-weighted mean based on the surface of each radiance pixel within a 200-m buffer around plot center. |
| **Noise intensity** | dB | For each plot, four measurements were taken at different locations within the plot and repeated on four separate sampling dates in spring 2023 (≥ 1 | (Schillé et al. 2025) | NA | Average of the 16 measurements within a 200-m buffer around plot center. |

| | | | | | |
|---|---|---|---|---|---|
| | | week apart), on different weekdays and times of day, with no fixed sequence to avoid confounding effects. | | | |
| **Surface temperature** | °C | Derived from 32-band thermal infrared imagery. Aerial surveys were conducted during daytime hours when impervious materials typically reach peak temperature (12:00–15:00), between August 3 and September 3, 2016. | (Ville de Montréal 2016) | 2 m spatial resolution | Area-weighted mean based on the surface of each thermography pixel within a 200-m buffer around plot center |
